# The PERL toolkit: Using sand flies to identify reservoirs of leishmaniasis

**DOI:** 10.64898/2026.06.27.734966

**Authors:** Eva Iniguez, Patrick Huffcutt, Tiago Donatelli Serafim, Pedro Cecilio, Serena Doh, Aaron Pugh, Johannes Doehl, Claudio Meneses, Ben Lambert, Jesus G. Valenzuela, Shaden Kamhawi

## Abstract

Animal reservoirs remain unknown in emerging and most endemic leishmaniasis foci hindering control efforts. Here, we developed a field-applicable PERL (Phlebotomines Establish Reservoirs of *Leishmania*) toolkit using individual blood fed sand flies (IBF) to identify leishmaniasis reservoirs. Using IBF, we optimized DNA and RNA co-extraction and parasite detection of ≥1 parasite/s by kDNA qPCR and *ssu rRNA* RT-qPCR. We then screened IBF for expression of two parasite genes, *sherp* and *HPB*, identified by RNAseq as having low-to-absent expression in IBF given a first *Leishmania donovani*-infected blood meal (IBF-iBM1) and high expression in specimens provided subsequent uninfected blood meals (IBF-BMS^+^). Linear discriminant analysis of target gene expression classified iBM1 parasites with a predictive accuracy of ∼87% and ∼82% in membrane- or naturally-fed on hamsters sand flies, respectively. By determining the blood source in specimens determined as IBF-iBM1, the PERL toolkit provides an innovative and practical approach to identification of leishmaniasis reservoirs.

## Introduction

*Leishmania*sis is a vector-borne disease caused by protozoan *Leishmania* parasites and transmitted by bites of female phlebotomine sand flies during blood feeding. The most prevalent clinical forms are cutaneous (CL) and visceral (VL) leishmaniasis. CL is generally self-healing and affects approximately a million people annually, while VL is responsible for up to 90,000 cases per year and is 95% fatal without treatment (1). *Leishmania*sis is mostly a zoonosis, involving animals that remain mostly unknown or unconfirmed. Only *Leishmania donovani* is thought to cause anthroponotic visceral leishmaniasis (AVL) (2) though recent studies suggest the likely presence of animal reservoirs (3, 4). These unidentified infection reservoirs persist as one of the most significant knowledge gaps in leishmaniasis foci worldwide, hindering development of efficacious and targeted control strategies to interrupt disease transmission. Lack of effective control of zoonotic foci also promotes expansion of leishmaniasis to new regions driven by factors such as climate and environmental changes, and population motility and displacement (5–10).

Xenodiagnosis is currently the most direct approach to determine the infectiousness of a *Leishmania*-infected host to sand flies and to investigate the role of suspected reservoirs (3, 11). However, this approach is laborious, costly, and selective, making it ineffective and infeasible for establishing the identity of unknown mammalian reservoir hosts in a field-setting. Other molecular biology-based methods are used to screen sand flies for infection or for identification of blood meal sources. However, they lack the sensitivity to distinguish initial parasite stages picked up from a reservoir animal from those residing in the gut during a subsequent blood meal by an infected vector (12, 13). Sand flies are promiscuous feeders that take blood meals every 5-6 days to support egg development (14). After a sand fly feeds on a disease reservoir, *Leishmania* begins a developmental cycle that spans multiple blood meals and multiple intermediate stages prior to maturation and transmission of infectious metacyclic promastigotes (14, 15). As a result, infected blood fed females collected in disease foci will contain distinct developmental stages of *Leishmania* parasites depending on the maturity of the infection. If we can distinguish *Leishmania* forms that are present in the first infected blood meal (iBM1) from those residing in a blood fed midgut during subsequent blood meals (BMS^+^), we can use sand flies to identify disease reservoirs in emerging or established leishmaniasis foci.

In this study, we developed an innovative field-applicable “Phlebotomines Establish Reservoirs of *Leishmania*” or PERL toolkit that focuses on in-depth sequential molecular analyses of DNA and RNA co-extracted from individual blood fed sand flies (IBF). Naturally, and without manipulation, this toolkit is tailored to point to leishmaniasis reservoirs by looking beyond an infection with *Leishmania* to specific parasite stages developing in the midgut in the context of the sand fly recurrent blood feeding behavior.

## Methods

### Animals

Male Golden Syrian hamsters (4-6 weeks old; Charles River Laboratories) and female BALB/c mice (3-6 weeks old; Jackson Laboratories) were maintained at the NIAID Twinbrook animal facility. Experiments were approved by the NIAID Animal Care and Use Committee (protocol LMVR4E) and conducted in accordance with NIH animal research guidelines.

### Parasites

*Leishmania donovani* (MHOM/SD/62/1S) was maintained by serial passage in Golden Syrian hamsters. Tissue amastigotes were harvested as described (16) and used to spike blood for sand fly infection or cultured to generate standard curves for qPCR. Parasites were cultured at 26°C in Schneider’s insect medium containing 20% heat-inactivated fetal bovine serum, 2 mM L-glutamine, and penicillin/streptomycin.

### Animal infections

Hamsters were infected intravenously with 10^7^ purified *L. donovani* metacyclic promastigotes. At 6-10 months after infection, symptomatic animals were exposed to sand flies or used to harvest splenic amastigotes (16).

### Sand fly infections and subsequent blood meals

Female *Lutzomyia longipalpis* (Jacobina strain; 4-6 days old) sand flies were infected by membrane feeding on animal blood containing 5 x 10^6^ tissue amastigotes/mL (15) or by feeding on an *L. donovani*-infected hamster (16). Blood-fed flies were maintained at 26°C with 30% sucrose. After the first infected blood meal (iBM1), flies received uninfected mouse blood meals at days 5-6 (BM2) and 10-12 (BM3). Midguts were collected at ≤2-48 hours after feeding; infection status and maturity were assessed as described (15).

### Preservation of individual blood fed sand flies on Whatman 903 Protein Saver Cards

After iBM1, BM2, or BM3, individual blood-fed midguts collected at ≤2-48 hours were spread onto Whatman 903 Protein Saver cards (Cytiva) (17), air-dried overnight, and stored individually at room temperature for up to 9 months. Dissection tools were decontaminated with 10% bleach between samples as described (12).

### DNA and RNA co-extraction from individual blood fed sand flies

Dried midguts were recovered with a 3-mm biopsy punch and DNA and RNA co-extracted using the Quick-DNA/RNA Miniprep kit (Zymo). Samples were digested in DNA/RNA Shield, proteinase K, and PK Digestion Buffer for 2 hours at 56°C and processed according to the manufacturer’s tissue protocol, including DNase I treatment of RNA. Nucleic acids were eluted in 100 uL and stored at −80°C.

### Leishmania probe-based kDNA qPCR

*Leishmania* kinetoplast minicircle DNA was quantified by TaqMan qPCR using JW11/JW12 primers and probe as described (18). Reactions contained 40 ng DNA and were run in duplicate or triplicate on a CFX96 Touch system (Bio-Rad). Each plate included an uninfected midgut control and a 10-fold standard curve generated from dilutions of an uninfected blood-fed midgut spiked with 10^5^ *L. donovani* parasites as described (12).

### Bulk RNA sequencing in infected blood fed sand flies

Pools of 20 or 5 infected midguts were collected at 0, 8, 24, and 48 hours after iBM1 or BM3 (day 10), respectively. Midguts were preserved in DNA/RNA Shield and extracted using the RNeasy Mini Kit (Qiagen). Libraries were sequenced on an Illumina NovaSeq 6000 (Novogene).

Reads were aligned with HISAT2 to a concatenated reference combininb *Lu. Longipalpis* and *L. donovani* genomes. *L. donovani* gene-level raw counts were retained. Retained samples required ≥10,000 parasite-aligned fragments and ≥300 parasite genes at counts per million (CPM) ≥1; these criteria retained iBM1 at 48 hours and all BM3 timepoints (3 biological replicates each). Candidate transcripts were subsequently validated by RT-qPCR in independent membrane- and hamster-fed sand flies.

Differential expression was analyzed in R with DESeq2 (19) after removing genes with <10 summed counts. Size factors were estimated using the post-counts method, and log2 fold changes were shrunk with apeglm (20). iBM1 parasite populations were compared with BM3 populations; adjusted p values were calculated by Benjamini-Hochberg correction. Among significant genes (adjusted p<0.05), candidates were prioritized by fold change and expression pattern.

### *sherp* identification and copy-number assessment in L. donovani

Because *sherp* is unannotated in the *L. donovani* BPK282A1 reference genome, the locus was identified by tBLASTn using *L. major* and *L. infantum sherp* proteins and manually annotated before read quantification. BLAST analysis supported a single assembled *L. donovani* copy and unambiguous read assignment. Locus identification, architecture, and cross-species copy-number analyses are provided in Figure S1.

### Quantification of live parasites and expression of *sherp* and *HPB* by One-Step RT-qPCR

Live parasites were quantified by one-step RT-qPCR targeting the constitutive *Leishmania* 18S ribosomal RNA (*ssu rRNA*) gene using a 10-fold RNA standard curve (21). Primers, forward (5’-GCGGAGACGAGCGACAAT-3’) and reverse (5’-CGAGCCGCCGCTTATCTT-3’), were designed to target the small hydrophilic endo-reticulum associated protein (*sherp*) gene in *L. donovani* (21, 22). For this study, forward (5’-GATGAGGACTGCAACACCCA-3’) and reverse (5’-AACGCCACAGGGGTGATAAG-3’) primers were designed to amplify the hypothetical protein (*HPB*) gene mined from our RNAseq data. Reactions using Luna Universal One-Step RT-qPCR master mix (NEB) were run for 10 minutes at 55°C, 1 minute at 95°C, then 40 cycles of 95°C for 10 seconds and 60°C for 30 seconds. Each plate included an uninfected blood-fed midgut control. ΔCT for parasite genes was calculated relative to the mean CT of the uninfected fed-midgut control.

### PCR for blood meal identification

To determine persistence of host DNA, sand flies were membrane-fed on human or cow blood. Midguts were collected immediately and every 24 hours through day 6. DNA was amplified using vertebrate mitochondrial cytochrome b (cow) and cytochrome c oxidase I (human) primers (23, 24) under previously described conditions (12).

### Classifying iBM1 from BMS^+^parasite populations

RT-qPCR data from 2 membrane-fed and 3 hamster-fed experiments were analyzed separately using the same pipeline. Nonamplifying *HPB* and *sherp* reactions were assigned CT=40; values >40 were capped at 40 and converted to abundance scores (40-CT). Samples were classified as iBM1 or subsequent blood meals (BMS^+^).

Linear discriminant analysis (LDA) used HPB and *sherp* abundance scores as predictors. Each dataset was stratified into 70% training and 30% held-out test sets. Models were implemented in R with tidymodels/MASS and evaluated by accuracy, F-measure, precision, and recall; the workflow was implemented with the targets package.

### Statistical analysis

Mann-Whitney and Kruskal-Wallis tests were used for 2-group and multiple-group comparisons, respectively. Spearman correlation assessed agreement between microscopy and molecular parasite estimates. Analyses were performed in GraphPad Prism 10.4.1; p≤0.05 was considered significant.

## Results

### Co-extraction of DNA and RNA from individual IBF midguts

We first optimized DNA/RNA co-extraction from individual blood-fed (IBF) midguts infected by membrane feeding (mean 849.2 ng ±SD 246.1 and mean 856.6 ng ±SD 265.2, respectively) or feeding on a clinically ill hamster (mean 864.9 ng ±SD 242.5 and mean 1135.0 ng ±SD 317.8, respectively). Each gut was preserved on Whatman 903 Protein Saver cards (Figure S2A,B). Nucleic acids sufficient for multiple assays were recovered from samples collected ≤2-48 hours postinfection, including from cards stored at room temperature for up to 9 months.

### Comparative parasite load assessment in IBF

To assess parasite burden after the first infected blood meal (iBM1), and after subsequent blood meals (BM2 and BM3), we quantified *Leishmania* by kDNA qPCR and *ssu rRNA* RT-qPCR in membrane- and hamster-fed IBF (Figure 1A-C; Figure S2C,D). Parasite estimates were substantially higher after membrane feeding than natural feeding at iBM1 (3.15-log by kDNA and 1.71-log by *ssu rRNA*) and increased after subsequent blood meals, particularly in hamster-fed flies (Figure 1B,C). Median kDNA estimates increased from 42,867 or 30 parasites at iBM1 to 562,579 or 977 at BM2 and 378,966 or 44,108 at BM3 for membrane- or hamster-fed IBF, respectively; corresponding *ssu rRNA* estimates were 1938 or 38, 105, 597 or 12,192, and 44,329 or 32,602 parasites for iBM1, BM2 and BM3, respectively (Figure 1B,C).

**Figure 1.**
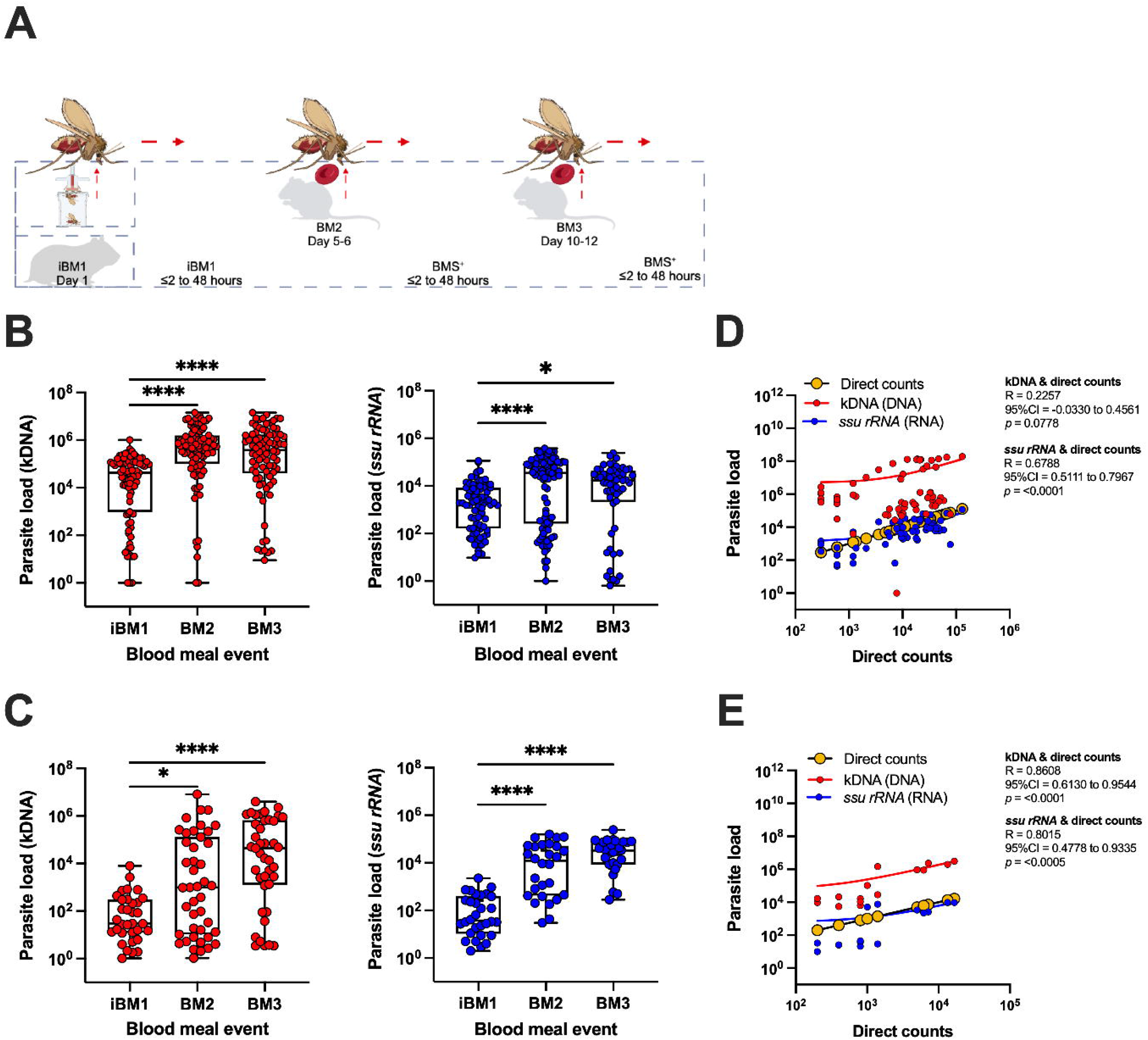
Establishing accurate quantification of *Leishmania* parasites from individual blood fed sand flies. (A) Experimental timeline. Sand flies were infected by membrane feeding on blood spiked with *L. donovani* parasites or on a clinically ill hamster. Sand flies were provided subsequent uninfected blood meals from mice at days 5-6 (BM2) and 10-12 post-infection (BM3). Individual fed midguts were collected from ≤2 to 48 hours after each blood meal event. (B,C) Parasites were molecularly quantified using a probe-based qPCR targeting kinetoplast DNA (kDNA, red circles), or RT-qPCR targeting the constitutively expressed parasite gene, *ssu rRNA* (blue circles), for sand flies fed on a membrane (B) or a clinically ill hamster (C). Cumulative data of two independent experiments are shown. (D,E) Parasite quantification by kDNA qPCR (DNA) and *ssu rRNA* RT-qPCR (RNA) was correlated with direct parasite counts by microscopy for a subset of samples from membrane-fed (D) or hamster-fed (E) sand flies. A standard curve was included in each plate to calculate parasite load per sample. Bar, mean ±95% CI. Kruskal Wallis test. *Spearman* correlation and coefficient of variation (R) with ±95% CI are provided for two independent experiments (D), or one experiment (E). (B,C) Each data point, mean of an IBF midgut ran in duplicate. iBM1, first infected blood meal; BM2, second uninfected blood meal; BM3, third uninfected blood meal; BMS^+^, subsequent blood meals. A p value of ≤0.05 was considered significant, ^∗^p < 0.05, ^∗∗∗∗^p < 0.0001. Created in BioRender. Iniguez, E. (2026) https://BioRender.com/lx0la30. Part of the Illustrations from NIAID Visual & Medical Arts, BioArt Source (bioart.niaid.nih.gov/bioart/000458) (A).

We compared molecular estimates with direct microscopy counts in a subset of iBM1 and BMS+ samples (Figure 1D,E; Figure S3A,B). In membrane-fed flies, microscopy counts correlated strongly with *ssu rRNA* (R=0.7, p<0.0001) but not kDNA (R=0.2, p=0.08). In hamster-fed flies, microscopy correlated with both *ssu rRNA* (R=0.8, p<0.0005) and kDNA (R=0.9, p<0.0001). Thus, kDNA may overestimate parasite burden after membrane feeding, whereas both assays accurately reflect parasite burden after natural feeding. Notably, *ssu rRNA* counts live parasites.

### *sherp* and *HPB* genes distinguish iBM1 from BMS^+^ parasite populations

We searched for transcripts that distinguish parasite populations present in iBM1 (iBM1pp) from those occurring in subsequent blood meals (BMS^+^pp). Candidate amastigote-associated genes, including amastin, DERA, LinJ, and A2, were expressed in BMS^+^pp and did not specifically identify iBM1pp (Figure S4A). In contrast, *sherp*, a metacyclic stage-enriched gene not previously annotated in the *L. donovani* reference genome (Figure S1), was not expressed by iBM1pp but similarly expressed by BM2pp and BM3pp (Figure S4B).

Next, we used bulk RNA sequencing to identify additional targets (Table 2). Consistent with targeted screening, amastin and A2 family genes were expressed at similar or higher levels in BM3pp than iBM1pp while *sherp* was upregulated in BM3pp (Figure S5A,B). We also identified a late-stage-enriched hypothetical protein, *HPB* (LdBPK_262710.1), whose expression was significantly lower in iBM1pp than BM3pp (adjusted p=1.1×10^-13^) and exceeded *sherp* expression across the tested BM3 timepoints (Figure S5B-D). Additionally, *HPB* showed high nucleotide identity with orthologs in clinically important *Leishmania* species (91.8%-99.8%; Table 3), similar to *sherp* and *ssu rRNA* (Table 3). Based on the above, we selected the late-stage-enriched *sherp* and *HPB* genes as targets for subsequent classification. Of note, candidate genes upregulated in iBM1pp were identified but they did not validate reliably by RT-qPCR despite extensive screening (Figure S6A,B),

**Table 1.** *sherp* locus architecture and copy number.

| Species<br>(assembly) | *sherp*<br>copies | Locus / arrangement | Gene IDs |
| --- | --- | --- | --- |
| *L. donovani*<br>(BPK282A1) | 1 | Chr 23 (Ld23_v01s1): single copy at<br>506,195–506,365 (–) | LdBPK_*sherp* (manually<br>added; unannotated in<br>TriTrypDB-68) |
| *L. infantum*<br>(JPCM5) | 2<br>(tandem) | Chr 23 (LinJ.23): identical copies ~4<br>kb apart, 510,745–510,918 and<br>514,920–515,093 (–) | LINF_230018650,<br>LINF_230018800<br>(100% identical) |

**Table 2.** Top ten transcripts differentially expressed at the initial infected blood meal (iBM1) versus BM3.

| Rank | Gene ID | Log<sub>2</sub><br>fold<br>change | adjusted<br>*p* value |
| --- | --- | --- | --- |
| 1 | LdBPK_310370.1 | 2.5 | 5.6E-17 |
| 2 | LdBPK_262710.1 (*HPB*) | 2.5 | 1.1E-13 |
| 3 | LdBPK_262600.1 | 2.5 | 1.6E-14 |
| 4 | LdBPK_040160.1 | 1.9 | 2.1E-11 |
| 5 | LdBPK_061320.1 | 1.5 | 4.7E-19 |
| 6 | LdBPK_170690.1 | 3.2 | 2.7E-11 |
| 7 | LdBPK_081190.1 | -1.1 | 8.3E-11 |
| 8 | LdBPK_323820.1 | 1.8 | 8.6E-11 |
| 9 | LdBPK_050900.1 | 2.4 | 1.7E-10 |
| 10 | LdBPK_190840.1 | 2.6 | 2.9E-10 |

iBM1, first infected blood meal; BM3, day 10.

**Table 3.** Percent identity of *HPB*, *sherp,* and *ssu rRNA* genes across *Leishmania* species. Alignment performed against available *Leishmania spp.* genomes.

| Gene | Description | GenBank Accession | Nucleotide Identity |
| --- | --- | --- | --- |
| ***HPB*** | *Leishmania donovani* hypothetical protein, unknown function (LDBPK_262710.1), partial mRNA | XM_003861807 | 100% |
|  | *Leishmania infantum* JPCM5 hypothetical protein, unknown function partial mRNA | XM_001470568 | 99.8% |
|  | *Leishmania major* strain Friedlin hypothetical protein partial mRNA | XM_001684223 | 94.6% |
|  | *Leishmania mexicana* MHOM/GT/2001/U1103 hypothetical protein, unknown function partial mRNA | XM_003876522 | 91.8% |
| ***Sherp*** | *Leishmania donovani* small hydrophilic endoplasmic reticulum-associated protein | Not annotated | - |
|  | *Leishmania infantum* JPCM5 small hydrophilic endoplasmic reticulum-associated protein | XM_003392466.1 | 100% |
|  | *Leishmania major* strain Friedlin small hydrophilic endoplasmic reticulum-associated protein (*sherp*) (*SHERP*), partial *mRNA.* | XM_001683388.1 | 90.2% |
|  | *Leishmania mexicana*<br>*MHOM/GT/2001/U1103 small hydrophilic*<br>*endoplasmic reticulum-associated protein*<br>*(sherp) partial mRNA* | XM_003875689.1 | 92.5% |
| **ssu rRNA** | *Leishmania donovani 18S ribosomal RNA*<br>*gene, complete sequence*<br>*(LdBPK_27rRNA6.1)* | GQ332356 | 99.7% |
|  | *Leishmania infantum JPCM5 18S ribosomal*<br>*RNA (SSU) RNA rRNA* | LINF_270031000-T1 | 99.7% |
|  | ***Leishmania major Friedlin 18S ribosomal***<br>***RNA (SSU) RNA*** | **XR_002460813** | **100%** |
|  | *Leishmania mexicana 18S ribosomal RNA*<br>*gene, complete sequence* | GQ332360 | 99.9% |

Bolded, reference gene.

### RT-qPCR validation of *sherp* and *HPB* expression in IBF

One-step RT-qPCR confirmed that *sherp* and *HPB* expression distinguished iBM1pp from BMS^+^pp in both membrane- and hamster-fed IBF (p<0.0001; Figure 2A,B). In membrane-fed flies, median ΔCT values increased from 8.4 for iBM1 to 14.0 (BM2) and 13.2 (BM3) for *sherp* and from 8.6 to 14.7 and 13.8 for *HPB* (Figure 2A). These patterns were reproduced in hamster-fed sand flies (Figure 2B), important to verify as parasite uptake at iBM1 was substantially lower than after membrane feeding. Moreover, separation between expression levels was more pronounced after natural feeding: median ΔCT increased from 0.0 at iBM1 to 10.4 and 13.1 at BM2 and BM3, respectively, for *sherp* and from 1.6 to 11.6 and 14.1 for *HPB* (Figure 2B).

**Figure 2.**
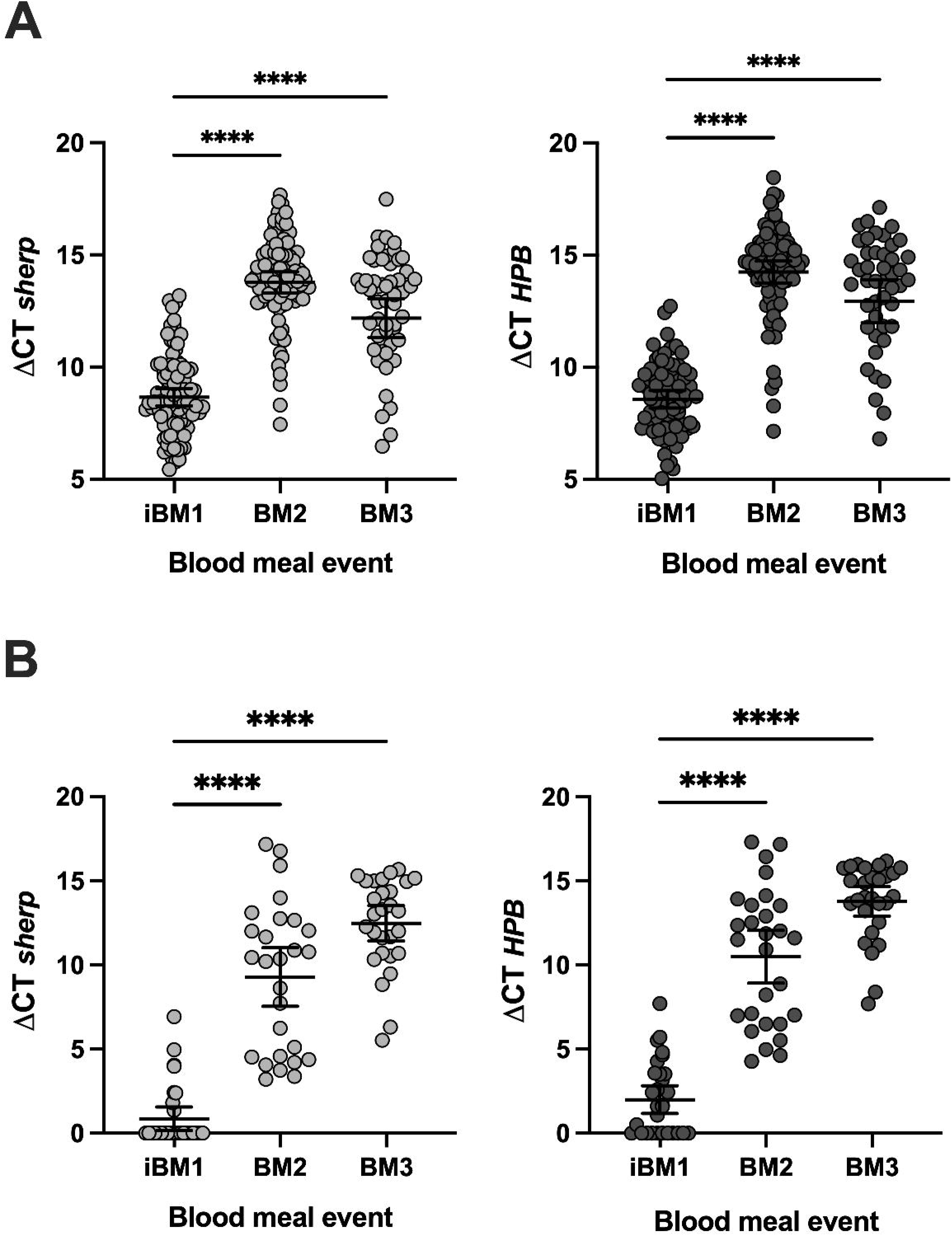
Stage-enriched expression of *Leishmania donovani* target genes distinguishes parasites within the first infected blood meal from those in subsequent blood meals in fed sand flies. (A,B) Expression of stage-enriched *sherp* or *HPB* at each blood meal in membrane-fed (A) or hamster-fed (B) sand flies. △CT for each gene was calculated by subtracting the mean CT value of a midgut fed on uninfected blood from the mean CT value of the infected sample; Bar, mean ± 95% CI are shown; Kruskal Wallis test. Cumulative data of two independent experiments are shown. Each data point represents the average of an IBF midgut ran in duplicate. A p value of ≤0.05 was considered significant, ^∗∗∗∗^p < 0.0001. iBM1, first infected blood meal on day 1; BM2, second uninfected blood meal on day 5-6; BM3, third uninfected blood meal on day 10-12. CT, cycle threshold.

### Blood meal persistence in blood fed sand flies

Because PERL works by linking iBM1pp to the vertebrate blood source, we fed sand flies on human and cow blood to demonstrate feasibility of blood meal identification and to determine how long host DNA remained detectable after feeding. In single fed specimens, human *COI* or Cow *cytb* DNA was detectable for up to 4 and three days, respectively (Figure 3).

**Figure 3:**
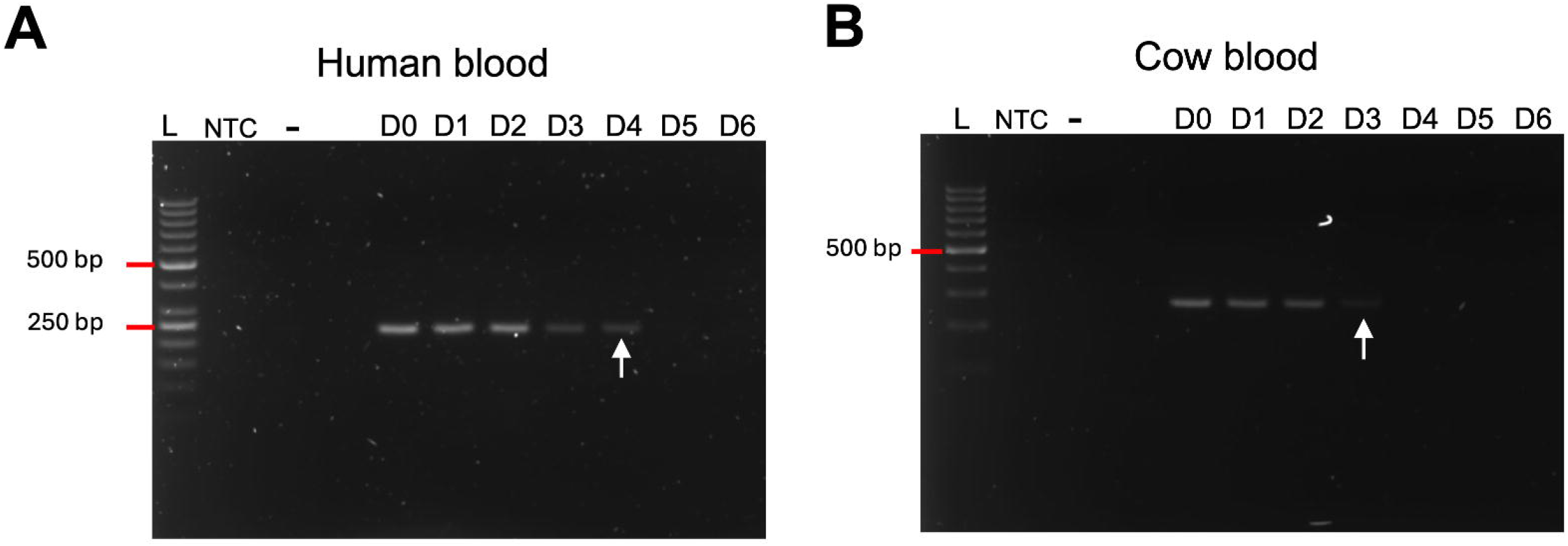
Detection of the blood meal source from fed sand flies. **(A,B)** PCR amplification of DNA from individual sand flies fed on a single host, human (A) or cow (B). Samples were collected right after feeding and daily thereafter up to day 6 post-feeding. Gel shows the distinct band sizes corresponding to cytochrome b (*cytb*) and cytochrome c oxidase subunit I (COI) of vertebrate hosts (cow, 263bp, *cyb*; human 228bp, COI). Negative controls: NTC, non-template control; -, DNA from an experimental sand fly fed on rabbit blood; L, ladder. Arrowhead, last day of detectable blood.

### Predictive accuracy of iBM1pp using a machine learning approach

Next, we used linear discriminant analysis to test whether combined *HPB* and *sherp* expression can distinguish iBM1pp from BMS^+^pp (Figure 4A-D). On held-out test data, classification accuracy was 87% for membrane-fed and 82% for hamster-fed sand flies (Table 4). Combined target performance was similar to or better than either target alone. These data support the feasibility of identifying iBM1pp using *HPB* and *sherp*.

**Figure 4.**
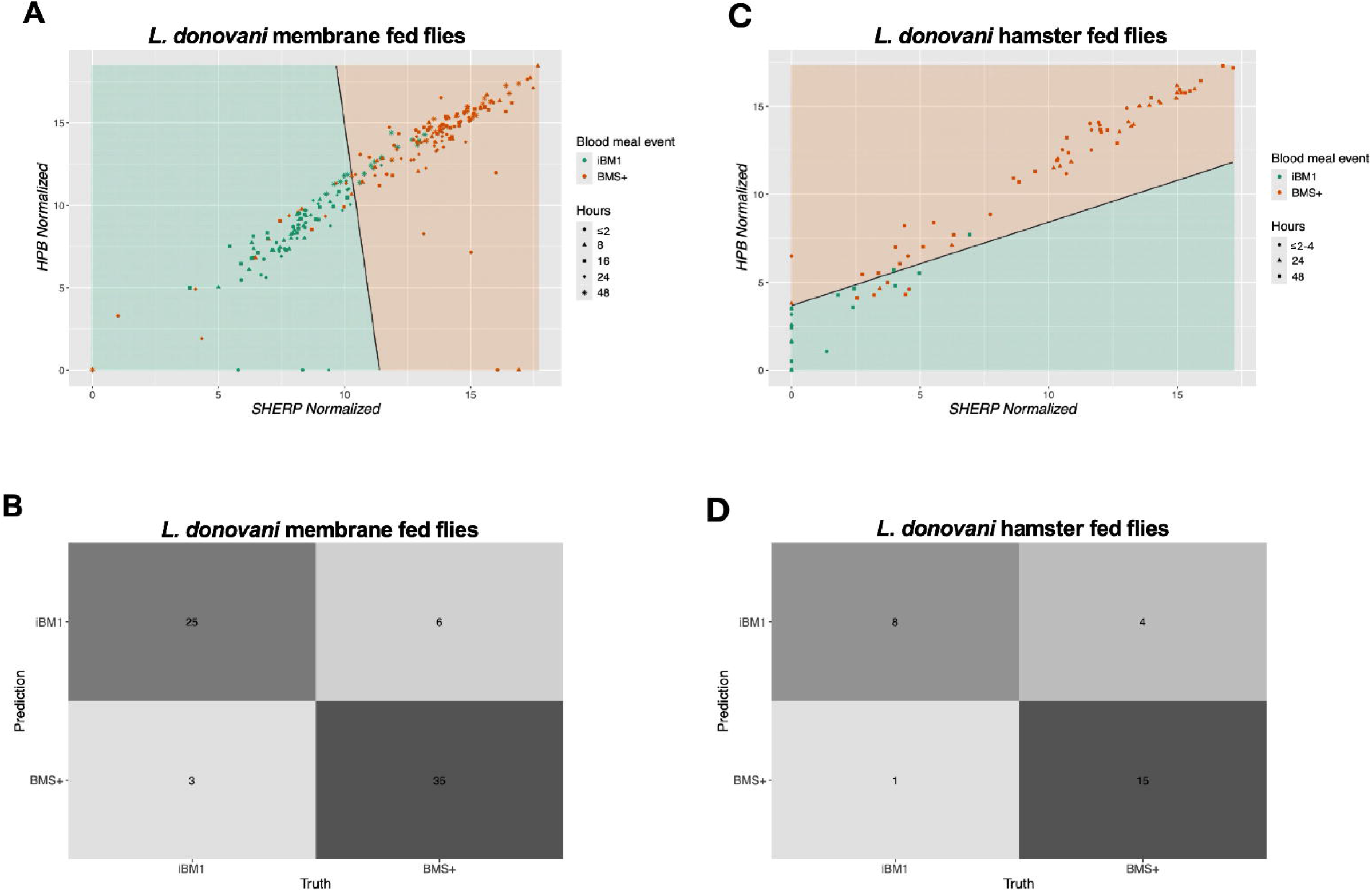
Target gene expression reliably differentiates the first *Leishmania donovani-* infected blood meal from subsequent blood meals in membrane and hamster fed sand flies. (A,B) Linear discriminant analysis using *HPB* and *sherp* as predictors. Black line, boundary determined by LDA fitted to the full dataset. (C,D) Confusion matrices determined by making predictions on an independent test set. iBM1, first infected blood meal. BMS^+^, subsequent blood meals.

**Table 4.** Test-set classification performance of the linear discriminant analysis (LDA) classifier of iBM1 from BMS^+^ in membrane-fed and hamster-fed sand flies. For the F-measure, precision and recall metrics, the BMS^+^ samples are treated as “positive” events.

| Infection | Predictors | Accuracy | F-measure | Precision | Recall |
| --- | --- | --- | --- | --- | --- |
| Artificial | *HPB+sherp* | 0.87 | 0.89 | 0.92 | 0.85 |
| Artificial | *HPB* | 0.81 | 0.84 | 0.87 | 0.80 |
| Artificial | *sherp* | 0.87 | 0.89 | 0.92 | 0.85 |
| Hamster | *HPB+sherp* | 0.82 | 0.86 | 0.94 | 0.79 |
| Hamster | *HPB* | 0.79 | 0.82 | 0.93 | 0.74 |
| Hamster | *sherp* | 0.79 | 0.82 | 0.93 | 0.74 |

### The PERL toolkit

PERL integrates parasite-stage classification with blood-meal identification. In Figure 5 we outline a working model based on *HPB* and *sherp* expression in the blood fed state, placing it in the context of the established life cycle of *Leishmania* parasites in the sand fly gut. After iBM1, ingested amastigotes transition to procyclic promastigotes within hours, and *sherp* and *HPB* expression remains absent or low (Figure 5A, upper panel). Expression of both targets starts to increase around 48 hours post iBM1 as parasites begin to exit the peritrophic matrix (PM) into the midgut environment (Figure 5A, lower panel). During BM2 and BM3^+^, promastigotes are present in the midgut, and we hypothesize that only some become trapped within the newly synthesized PM surrounding BMS^+^ (Figure 5B-C). Under these conditions, *sherp* and *HPB* expression is significantly higher than in iBM1pp. Thus, absent-to-low expression of *HPB* and *sherp* identifies parasites associated with the first infected blood meal, whose vertebrate DNA can then be analyzed to identify a potential reservoir host.

**Figure 5.**
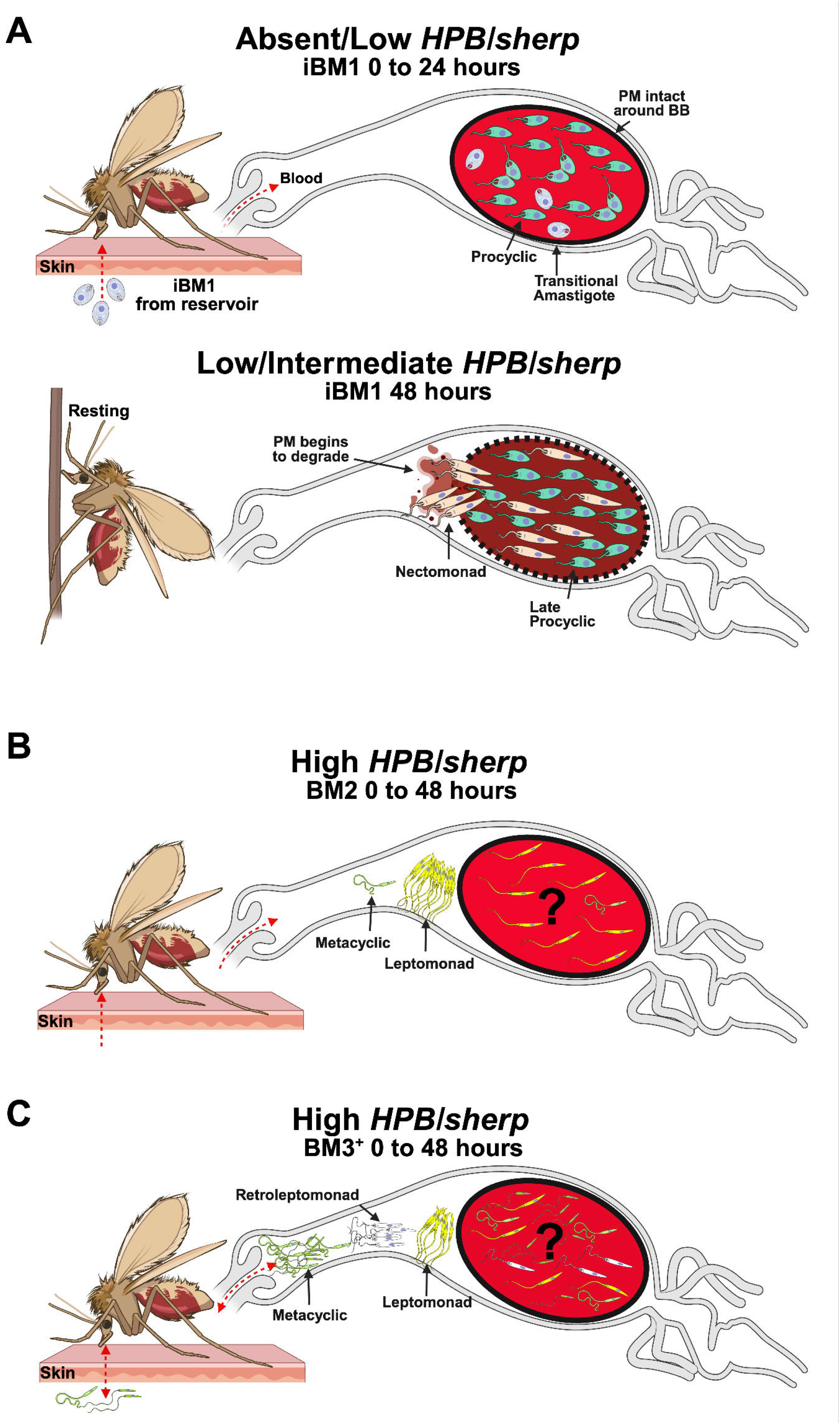
Identifying animal reservoirs by deploying a field-friendly sand fly toolkit in leishmaniasis foci. (A) During the first infected blood meal (iBM1) sand flies ingest amastigotes from an infected reservoir. In the sand fly midgut, amastigotes transition to procyclic promastigotes and divide by 24 hours. At this stage of the infection, the iBM1 parasite population (iBM1pp) is contained within the blood bolus (BB) and protected by the peritrophic matrix (PM). In this scenario *sherp* and *HPB* expression is absent or detected at very low levels (A, top). Around 48 hours after the initial infected blood meal, the PM begins to degrade as blood digestion progresses, permitting nectomonad promastigotes to exit into the lumen and attach to the midgut epithelium (A, bottom). At this timepoint, iBM1pp contains mainly nectomonad forms with few procyclic promastigotes and shows low to intermediate *sherp* and *HPB* expression (A, bottom). (B,C) The *Leishmania*-infected sand fly takes subsequent blood meals (BMS^+^) every 5-6 days. (B) During BM2, leptomonad promastigotes predominate and a few infectious metacyclic promastigotes may appear. (C) During BM3^+^, several parasite stages may be present, including leptomonad, retroleptomonad, and infectious metacyclic forms. In both BM2pp and BM3^+^pp, *sherp* and *HPB* expression is significantly higher than in iBM1pp (B,C). Upon imbibing a BM2 or BM3^+^, the location of parasites with respect to the PM remains unknown (?). The most likely scenario is that parasites may reside both inside and outside the BB (B,C). iBM1, BM2, and BM3^+^ depict a blood meal given on day 1 (iBM1), day 5-6 (BM2), or day 10-12 and after (BM3^+^). Created in BioRender. Iniguez, E. (2026) https://BioRender.com/hah62lk. Part of the Illustrations from NIAID Visual & Medical Arts, BioArt Source (bioart.niaid.nih.gov/bioart/290, 291, 294, 296, 458 and 461).

## Discussion

*Leishmania* species that are pathogenic to humans are mostly zoonotic in nature (25), yet the identity of animal reservoirs in many endemic and emerging disease foci remains in question (3, 4, 25), posing a serious challenge to control efforts. This is partly because adequate screening of animals, particularly if they are wild and diverse, is difficult and costly. In this study we introduce the field-friendly PERL toolkit comprising a progressive set of analyses that determines potential animal reservoirs of leishmaniasis from blood fed female sand flies. As such, PERL provides an additional and epidemiologically vital piece of information from field-captured *Leishmania*-infected blood fed sand flies (12, 26, 27).

PERL innovates by its ability to distinguish parasites developing in a first infected blood meal, taken by a sand fly from its reservoir host, from those present in subsequent blood meals, taken by infected females from prevalent animals. We hypothesize that our ability to distinguish iBM1pp from BMS^+^pp is related to the unique environment of the former which resides in a blood bolus taken from the reservoir host. Amastigotes taken up in the blood meal are contained by a PM that separates the parasites from the sand fly midgut. This condition is unique in that the developing parasite population has not yet been in contact with the midgut environment. Our data show that at 48h post-iBM1, which coincides with the initiation of PM degradation (28, 29), the parasites begin to upregulate *HPB* and *sherp* expression, potentially as an adaptation to the new midgut environment. During PM degradation, the parasites escape into the midgut lumen where they interact with the epithelium for the duration of the sand fly lifespan. This hypothesis is supported by the unexpected observation that *sherp*, a well-studied gene that has been closely linked to the development of infectious metacyclic promastigotes (22, 30–32), is highly expressed on day 6 post-infection when this stage is absent or negligible. Moreover, *sherp* expression remains constant as the infection matures, and seems to be more related to contact with the midgut environment than to the appearance of infectious metacyclic promastigotes. Notably, amastigote-associated genes such as the *Amastin* and *A2* genes were strongly expressed by promastigotes developing in both iBM1 and BMS^+^ sand flies. Together, these data indicate that *sherp* is not a reliable candidate for the detection of infectious sand flies and emphasize the relevance of conducting studies in vector sand flies to validate gene expression of candidates identified as stage-specific or stage-enriched under culture conditions.

Blood feeding is a prominent event that recurs throughout the life span of sand flies. If we consider that the sand fly takes a blood meal every 5-6 days (14), and that blood takes up to 3-4 days to be digested, then the parasites are mostly in contact with blood. Despite this intimate association between the blood fed state and *Leishmania*, fundamental information about the physical whereabouts of the parasites within the midgut following the intake of a subsequent bloodmeal and de novo synthesis of a PM remains unclear. We hypothesize that upon ingestion of blood, the newly synthesized PM may potentially trap a subset of parasites inside the new blood meal while others remain associated with the midgut lumen creating two subpopulations that are experiencing very distinct environments. This is supported by our data where iBM1pp and BMS^+^pp show differences in gene expression making the toolkit possible. Knowing that blood profoundly affects the development of *Leishmania* parasites (15, 33), establishing whether these two subpopulations occur and whether they are differentially reprogrammed would reveal novel and important aspects of host-parasite-vector interactions. For example, understanding the distribution of midgut-residing parasites in the context of recurrent blood meals may have potential implications for how hybrids, reported from natural isolates (34, 35), may form.

Throughout this study we investigated the expression of our target genes, *HPB* and *sherp*, in midgut-residing parasites after feeding on an artificial membrane or a sick hamster. As reported previously (15), our data established a lower iBM1 parasite count estimate for naturally compared to artificially fed sand flies, emphasizing the need to verify translational tools in a more natural system. The toolkit performed well, with a predictive accuracy above 80%, in both membrane- and hamster-fed groups demonstrating the feasibility of PERL in a natural setting.

In our controlled laboratory experiments, exposing the sand flies to a membrane feed or a sick animal takes a minimum of 1-2h, hence our earliest timepoint is indicated as ≤2h. Practically, investigators are unlikely to encounter amastigotes in sand flies from field captures unless a rare specimen is aspirated as it feeds on an infected animal. Even in this situation, the time to sorting and processing may add hours to the time of collection. This is relevant from a translational perspective where we expect that the vast majority of field collected blood fed sand flies will contain parasites that have been developing in the midgut for 8h or longer.

There are several limitations to our study. Our initial objective was to identify genes that were specifically upregulated in iBM1pp and BM3^+^pp to distinguish a specimen that fed on a reservoir animal from an infectious specimen, respectively. Unfortunately, though RNAseq of pooled sand flies revealed a few upregulated genes in iBM1 parasites, all failed validation in RT-qPCR of individual sand flies emphasizing the inadequacy of candidate selection based on RNAseq alone. Importantly, on a transcriptional level, BM2pp and BM3pp showed near identical expression of *HPB* and *sherp*. Future work is needed to mine better targets, not necessarily based on gene expression, that may be able to perform better at distinguishing iBM1pp, BM2pp, and BM3^+^pp thus reliably identifying infectious specimens in addition to reservoir animals. Further, the PERL toolkit depends on finding enough *Leishmania*-infected blood fed specimens. This requires knowledge and expertise in where and when to trap sand flies in a study focus, and comprehensive collections to gather enough blood fed infected specimens from ecotypes of interest.

This work puts forward an innovative idea of how we can use sand flies to find unknown leishmaniasis reservoirs in endemic or emerging disease foci. The PERL toolkit is versatile and can be deployed in any active leishmaniasis focus, including those in remote areas due to its field-friendly translational approach of preserving both DNA and RNA on filter paper. Considering that thousands of sand flies are captured and analyzed in numerous entomological studies of leishmaniasis foci, applying the PERL toolkit to collected specimens has the potential to be the most efficient way of finding unknown animal reservoirs.

## Contributions

E.I. and S.K. conceived the study, designed the experiments, analyzed data, supervised the project, and wrote the manuscript. E.I. and P.H. performed the majority of the experiments. P.H. contributed to experimental design and conducted data analysis. T.D.S. performed RNA-sequencing data analysis. P.C., S.D., A.P., J.D. and C.M. assisted with experiments. B.L. and J.G.V. contributed to data interpretation and modeling, scientific discussion, and manuscript writing. All authors reviewed, edited, and approved the final manuscript.

## Acknowledgments

We thank the Research Technologies Branch of the Rocky Mountain Laboratories at NIAID/NIH for design of primers specific for expression of sherp for *L. donovani* and the insectary students at the Twinbrook III insectary facility at the LMVR, NIAID/NIH. Illustrations from NIAID NIH BioArt Source (bioart.niaid.nih.gov/bioart). ChatGPT version 5.6 Luna was used solely to reduce the word count to meet journal requirements. The authors carefully reread and revised the reduced text ensuring its accuracy and completeness and take full responsibility for the content.

## Funding

This research was supported by the Intramural Research Program of the National Institutes of Health (NIH). The contributions of the NIH authors are considered Works of the United States Government. The findings and conclusions presented in this paper are those of the authors and do not necessarily reflect the views of the NIH or the U.S. Department of Health and Human Services.

## Data availability

All normalized and raw RNA seq data comparing BM1 vs BMS^+^ will be deposited at GitHub depository and available upon acceptance. All scripts used for data processing and analysis will be available at GitHub depository upon acceptance.

## Conflict of interest

The authors declare no conflict of interest.

## Notes

### Competing Interest Statement

The authors have declared no competing interest.

### Summary of Updates

The manuscript was shortened to approximately 3,500 words, and the figures were reordered accordingly. These revisions did not alter the scientific content, results, or conclusions of the manuscript.

