## Supplemental for "The PERL toolkit: Using sand flies to identify reservoirs of leishmaniasis"

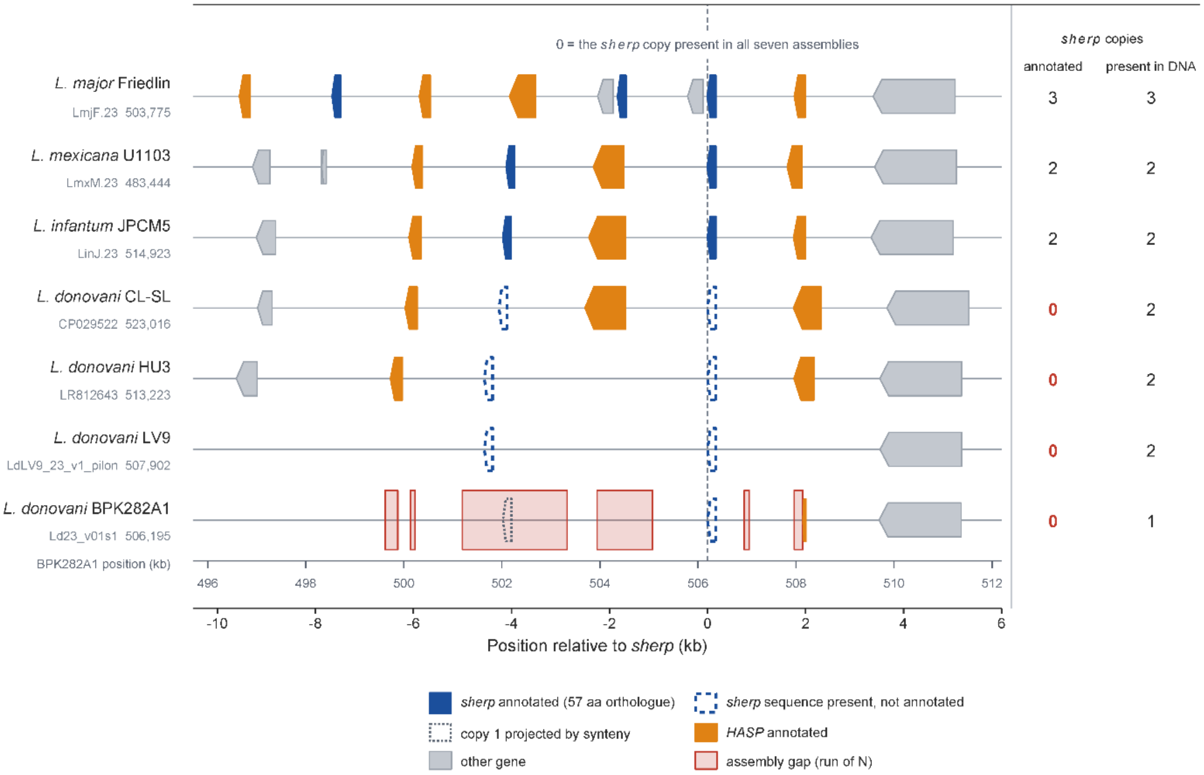


**Figure S1. Comparative architecture and annotation of the HASP/*sherp* locus across seven *Leishmania* genome assemblies**. Chromosome 23 locus structure is shown for *L. major* Friedlin, *L. mexicana* U1103, *L. infantum* JPCM5, and four *L. donovani* assemblies (CL-SL, HU3, LV9, and BPK282A1). The dashed vertical line at position 0 relative to *sherp* (x = 0) marks the *sherp* hit selected as the alignment anchor in each assembly; this anchor is recovered in all seven assemblies. Because each row is plotted in its own assembly coordinates, x = 0 has a different absolute contig or scaffold coordinate in each row, provided by the gray text beneath each taxon. The lower axis reports distance from this anchor. The upper secondary ruler gives absolute coordinates for the BPK282A1 reference assembly and applies only to the bottom row. Filled blue arrows mark *sherp* features in the TriTrypDB release 68 annotations (174-bp CDS including the TAA stop codon; predicted 57-amino-acid *sherp*); dashed blue arrows outline full-length matches to the 171-bp, stop-excluded *L. donovani* open reading frame recovered by ungapped seed-and-extend matching (no annotated feature of any type lies within 300 bp); orange arrows mark annotated HASP genes; gray arrows depict other annotated features; pale red rectangles indicate N-runs; and the dotted gray outline shows the expected BPK282A1 copy position projected by local synteny from *L. infantum* JPCM5. Arrowheads show transcriptional direction. The right-hand columns report the number of released *sherp* annotations and the number of full-length *sherp* sequence hits in each assembly. Thus, *sherp* sequence is recovered from all four *L. donovani* assemblies but is absent from the corresponding TriTrypDB release 68 annotations. In the BPK282A1 reference assembly, six N-runs interrupt the displayed interval. Local syntenic projection from *L. infantum* JPCM5 places the expected 174-bp position of the unresolved copy entirely within the largest N-run (2,140 placeholder Ns). N-run width does not establish the length or content of missing sequence. The three-copy count for *L. major* is specific to the reference LmjF.23 assembly. The Friedlin2021 reassembly of the same strain contains two copies.

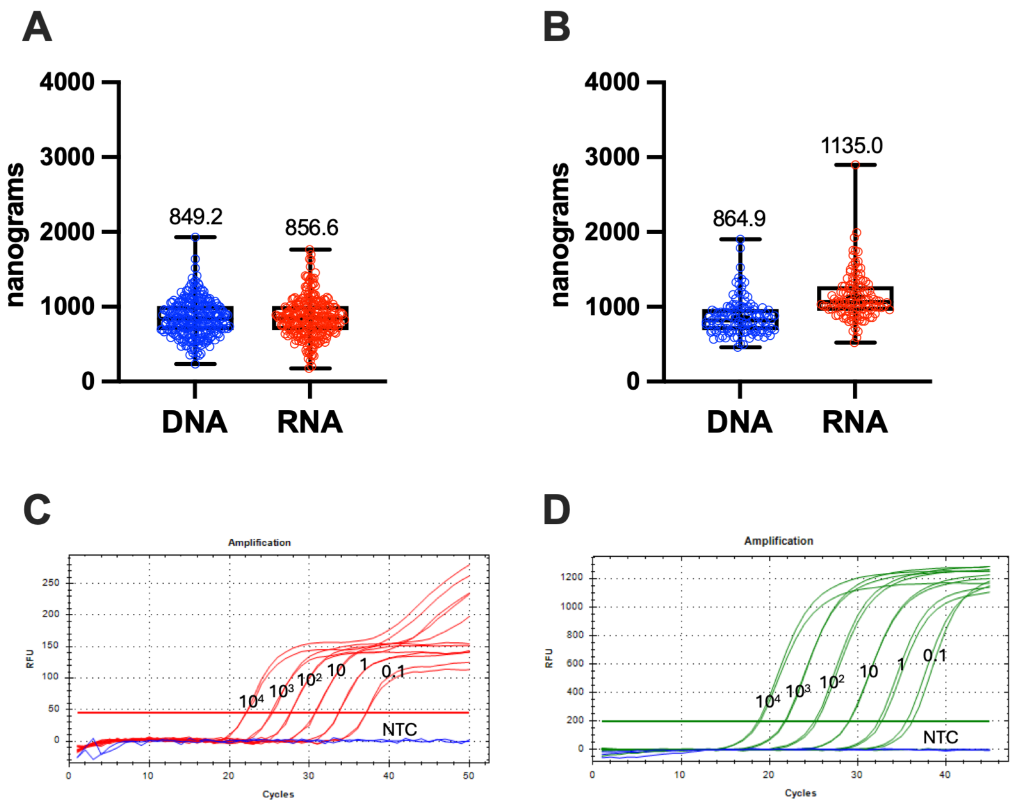


**Figure S2. Co-extraction of DNA and RNA from individual blood fed sand flies and generation of standard curves.** (A,B) Quantity of DNA and RNA extracted from *Leishmania*-infected midguts collected at ≤2 to 48 hours post-infection and preserved on Whatman 903 Protein Saver cards. Sand flies were either fed on a membrane (A) or a clinically ill hamster (B). Nucleic acids were extracted from samples stored at room temperature for up to 9 months. Bar, mean ± 95% CI. Each data point, an individual blood fed sand fly midgut quantified by nanodrop. (C,D) Representative amplification plots of a standard curve targeting kDNA by qPCR (C) or *ssu rRNA* by RT-qPCR (D). A relative fluorescent unit threshold was set for all the plates at 45 for (C) and 200 for (D). NTC, non-template control (blue).

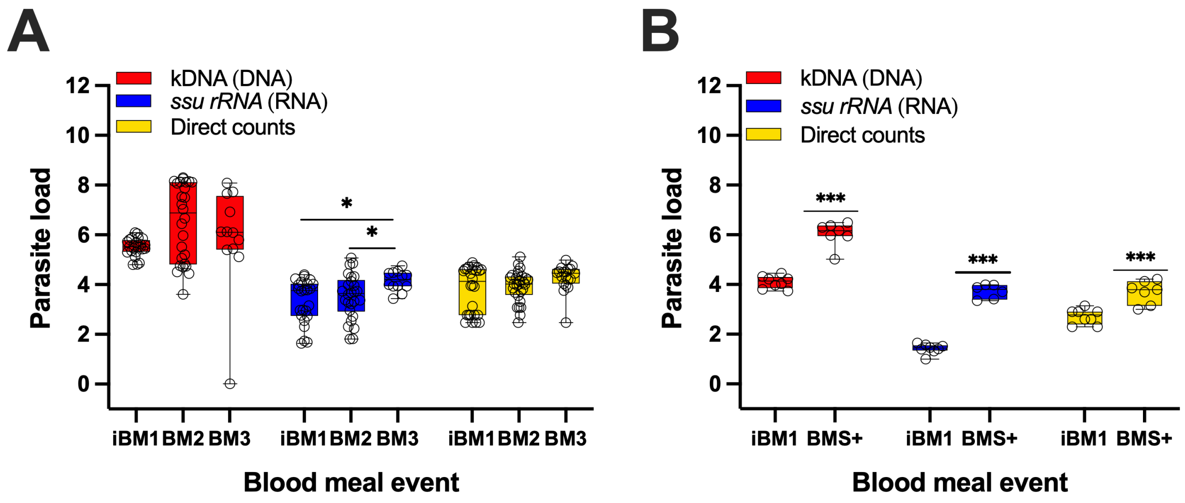


**Figure S3. Validation of molecular targets for parasite quantification by comparison with microscopy counts in individual fed sand flies.** (A,B) Parasites were molecularly quantified using a probe-based qPCR targeting kinetoplast DNA (kDNA), or RT-qPCR targeting the constitutively expressed parasite gene, *ssu rRNA*, and compared to direct counts by microscopy for sand flies fed on a membrane (A) or a clinically ill hamster (B). A standard curve was included in each plate to calculate the parasite load per sample. Cumulative data of two independent experiments (A), or one experiment (B). Bar, mean ±95% CI. Mann-Whitney (A) or Kruskal Wallis (B) test. A p value of ≤0.05 was considered significant, $^{***}p < 0.001$, $^{*}p < 0.05$. For kDNA and *ssu rRNA*, each data point represents the mean of an IBF midgut ran in duplicate. iBM1, first infected blood meal; BM2, second uninfected blood meal on day 5-6; BM3, third uninfected blood meal on day 10-12; $BMS^{+}$, subsequent uninfected blood meal on day 10.

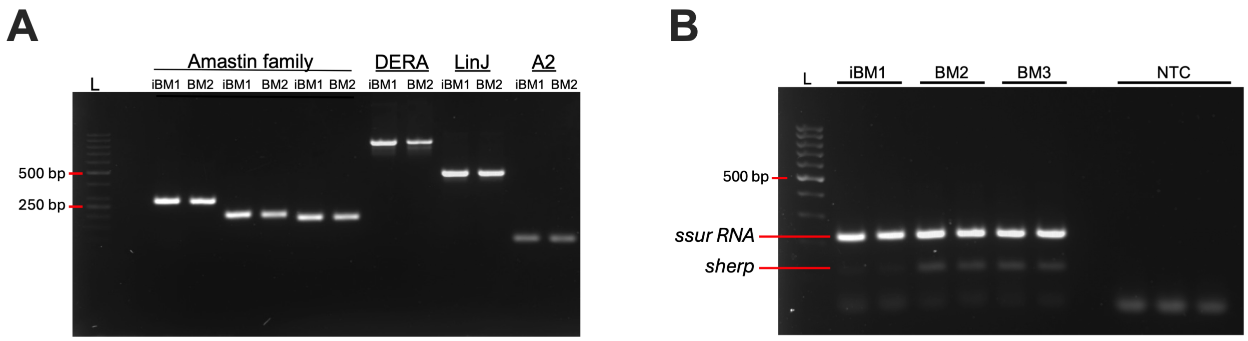


**Figure S4: Previously characterized stage-specific genes are expressed across other parasite stages in the sand fly midgut.** (A,B) Screening of amastigote genes (A), and *sherp* (B) in individual membrane fed sand flies by RT-PCR. Housekeeping gene *ssu rRNA* was screened in parallel (B). Representative samples from two independent experiments are shown; NTC, non-template control; L, ladder; iBM1, infected blood meal on day 1; BM2, second uninfected blood meal on day 6; BM3, third uninfected blood meal on day 12.

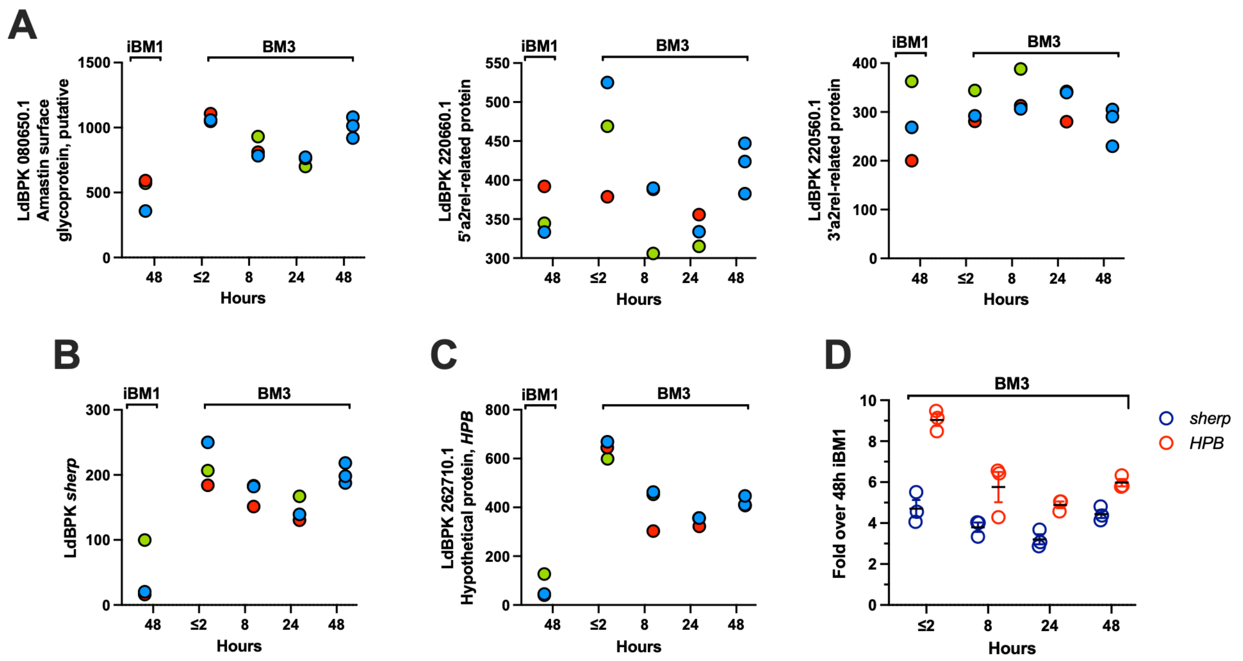


**Figure S5. Discovery of *HPB,* a novel gene that distinguishes parasites residing in the sand fly midgut during the first infected blood meal from those in subsequent blood meals.** (A-C) Bulk RNA-seq expression of the amastin gene and two members of the A2 gene family (A), *sherp* (B), and *HPB,* a novel target gene coding for a hypothetical protein (LdBPK_262710.1) (C). (D) Comparison of expression levels between *HPB* and *sherp*. Pools of 20 (iBM1), or 5 (BM3) fed midguts were collected per time point at 48 hours after iBM1, and ≤2, 8, 16, and 48 hours after BM3. Data from 3 independent experiments are shown. Graphs show the CPM (counts per million) representing the average gene expression level per experiment (A-C) or fold-change over 48h iBM1 (D). iBM1, first infected blood meal; BM3, a subsequent uninfected blood meal given on day 10 post infection.

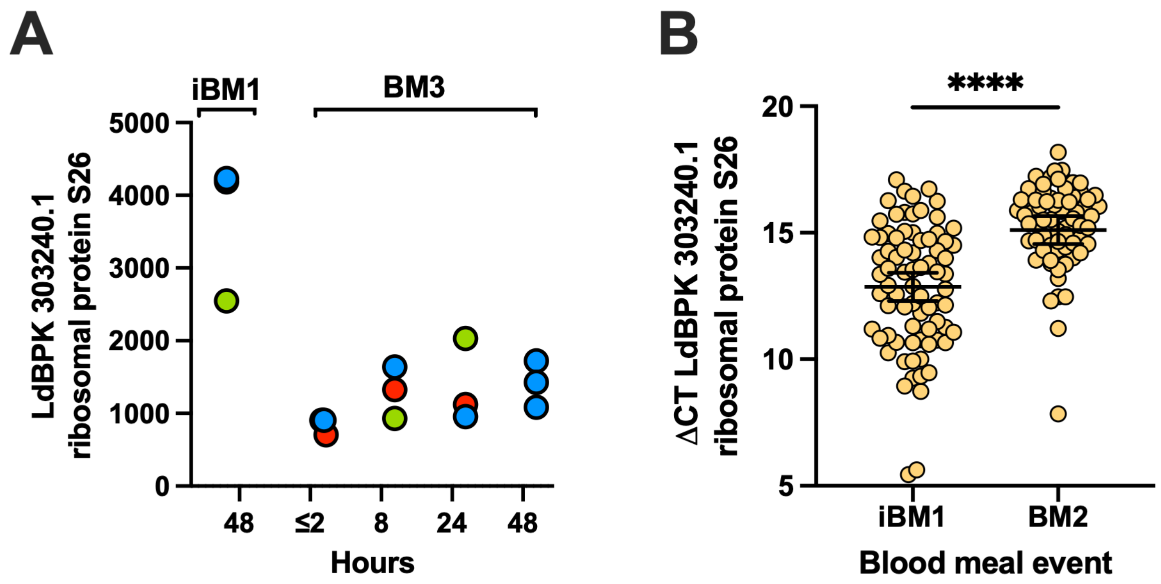


**Figure S6. Validation of the most upregulated gene in the first infected blood meal compared to subsequent blood meals**. (A) Bulk RNA-seq expression of gene LdBPK_303240.1, a ribosomal protein S26 upregulated in the iBM1pp. Pools of 20 (iBM1), or 5 ($BMS^{+}$) fed midguts were collected per time point at 48 hours after iBM1, and ≤2, 8, 16 and 48 hours after BM3. Data from 3 independent experiments are shown. Graphs show the CPM (counts per million) representing the average gene expression level per experiment. (B). Gene validation by qRT-PCR. ∆CT for each gene was calculated by subtracting the mean CT value of a midgut fed on uninfected blood from the mean CT value of the infected sample; Bar, mean ± 95% CI are shown; Mann-Whitney test. Cumulative data of two independent experiments are shown. Each data point represents the mean of an IBF midgut ran in duplicate. iBM1, first infected blood meal; BM2, a second uninfected blood meal on day 6; BM3, a subsequent uninfected blood meal on day 10.
